# Metabolic Heterogeneity of Muscle Stem Cells is Controlled by the Myofiber Niche

**DOI:** 10.64898/2026.09.04.749367

**Authors:** Clovis Chabert, Marie Quétin, Laurence A. Neff, Alessandro Bonavoglia, Olivier M. Dorchies, Philippos Mourikis, Marianne Gervais, Frederic Relaix, Alexandre Prola

## Abstract

Metabolic pathways support biomass synthesis and provide substrates for epigenetic modifications of histones and DNA. These findings are particularly relevant for adult stem cells, where metabolic changes significantly impact functional behavior. However, assessing the metabolism of these cells in their native microenvironment is challenging as this typically necessitates dissociation prior to metabolic analysis. In this study, we focus on muscle stem cells (MuSCs) and show that removing these cells from the niche significantly alters their metabolic profile. To overcome this issue, we developed a novel enzymatic staining method for *in situ* metabolic profiling at a single-cell resolution. This approach reveals unexpected metabolic heterogeneity of MuSCs, identifying oxidative and glycolytic subsets, and demonstrates that their metabolism is directly modulated by adjacent myofibers. Accordingly, perturbing myofiber metabolism remodels the MuSC niche and drives metabolic adaptation in MuSCs. Finally, we define the kinetics by which myofiber-dependent metabolic regulation of MuSCs is re-established during muscle regeneration, thereby revealing how niche-imposed metabolic cues shape stem cell identity *in vivo*.

**Teaser:** A new *in situ* approach reveals how the muscle niche controls stem cell metabolism.

## Introduction

Cells continuously adapt their metabolism to environmental conditions, particularly fluctuations in nutrient and oxygen availability. This metabolic flexibility is essential, as even a brief interruption in ATP resynthesis would rapidly deplete cellular ATP stores. Such flexibility relies on two processes operating on distinct timescales. First, rapid metabolic adjustments occur through transcription-independent mechanisms that modulate enzymatic activity, including allosteric regulation, enzyme clustering, and post-translational modifications (*1–3*). Second, long-term metabolic adaptations involve the activation of transcriptional programs that increase the expression of specific metabolic enzymes, to promote sustained shifts in pathway usage (for review see (*4*). The substrate preference for glucose, fatty acids or amino acids is of crucial importance since the production of secondary metabolites associated with them induces specific genetic programs through the regulation of (epi)genetic reactions (*4–6*). Consequently, metabolic adaptation to environmental fluctuations plays a crucial role in determining cell fate. However, this role remains poorly understood due to the technical challenges of studying cells within their native microenvironment. Adult stem cells offer an ideal model system to investigate how environmental cues shape metabolism and influence fate decisions, an area of significant therapeutic potential. However, our understanding remains limited, as most current insights into stem cell metabolism are derived from isolated or cultured cells, where the influence of the native microenvironment, the so-called niche, is largely lost and insufficiently explored.

Adult muscle stem cells (MuSCs), also known as satellite cells, confer high regenerative capacities to skeletal muscle. This tissue is indeed able to adapt its mass in response to functional needs and to regenerate in response to injuries. MuSCs are normally quiescent, a state characterized by the expression of the transcription factor paired box 7 (PAX7) (*7*). In response to muscle injury, MuSCs are activated, enter into a proliferating state, and start expressing myogenic differentiation factor 1 (MYOD1). A subset of cells then commits to differentiation, loses PAX7 expression and starts expressing myogenin (MYOG). Differentiated cells then fuse with differentiating myogenic cells to form new myofibers or with pre-existing myofibers to sustain muscle growth or repair. Importantly, a portion of activated MuSCs does not differentiate but instead downregulates myogenic gene expression and exits the cell cycle to re-enter into quiescence to self-renew the PAX7^+^ MuSC pool. During the last decades, alterations of MuSC function and reduced muscle regenerative capacities have been described in numerous pathological conditions, which irremediably contribute to muscle dysfunction and loss. The interaction of MuSCs with their environment appears critical in this context since dysfunctional MuSCs in aged or dystrophic mouse models regain function when grafted into young healthy hosts (*8*, *9*). However, the nature of this environmental regulation of MuSC function remains to be determined.

Growing evidence suggests that changes in the myogenic state of MuSCs are associated with metabolic shifts (*10–20*). It has been proposed that targeting MuSC metabolism could be an attractive approach to control their state and thus their regenerative capacities. This hypothesis is supported by genetic or pharmacological studies demonstrating that metabolic pathways are essential for regulating MuSC function. As an example, lactate exposure enhances myoblast differentiation through epigenetic mechanisms (*21*). However, the metabolic identity and dynamics of MuSCs in vivo, as well as the influence of their microenvironment on their metabolic state, remain poorly understood, since most available data have been obtained from dissociated cells or from cells studied in culture. To overcome this limitation, we developed an innovative method that enables *in situ* metabolic measurements at single-cell resolution, allowing direct investigation of MuSC metabolism within its native niche. This approach revealed an unexpected metabolic heterogeneity among MuSCs and showed that their metabolic state is strongly influenced by the surrounding myofibers. Finally, we define the kinetics by which myofiber-dependent metabolic regulation of MuSCs is re-established during muscle regeneration, revealing how niche-imposed metabolic cues shape stem cell fate *in vivo*.

## Results

To evaluate whether isolating MuSCs from their microenvironment disturbs their metabolism, we first analyzed previously published RNA-seq dataset to examine the expression of metabolic genes in MuSCs fixed with formaldehyde at various time points during the standard dissociation procedure (*22*). We observed a global decrease in the expression of genes involved in fatty acid β-oxidation and mitochondrial respiration as early as 30 to 60 minutes after initiation of the isolation process of MuSCs from their microenvironment (Fig. S1a). These changes were associated with an increase in the expression of genes involved in glycolysis and amino acid catabolism. Since these metabolic changes might be related to the transition of stem cell state (i.e., the early activation of MuSCs reported by Machado et al. 2017 during the isolation step), we repeated this experiment using both MuSCs and endothelial cells (ECs) isolated from Tg:Pax7-nGFP; VE-Cadherin-tomato mice. This mouse model allowed us to compare the impact of dissociation on metabolic genes (a selected list of 184 genes) in FACS-sorted stem cells (MuSCs, Pax7-nGFP^+^) or non-stem endothelial cells (EC, VE-CADHERIN^+^, Fig. 1a). In response to isolation, 78% of genes involved in metabolic pathways were commonly up- or downregulated in both MuSCs and ECs. This resulted in a strong correlation in dissociation-induced log2FC between MuSCs and ECs (Fig. 1b). Among the commonly downregulated genes, we found genes involved in fatty acid oxidation (Fig. 1c), while among the commonly upregulated genes, we found genes involved in amino acid catabolism and glycolysis (Fig. 1d-e). The comparable transcriptomic responses in ECs suggest that the effects of isolation reflect a general cellular stress response rather than a feature specific to adult stem cells.

**Figure 1.**
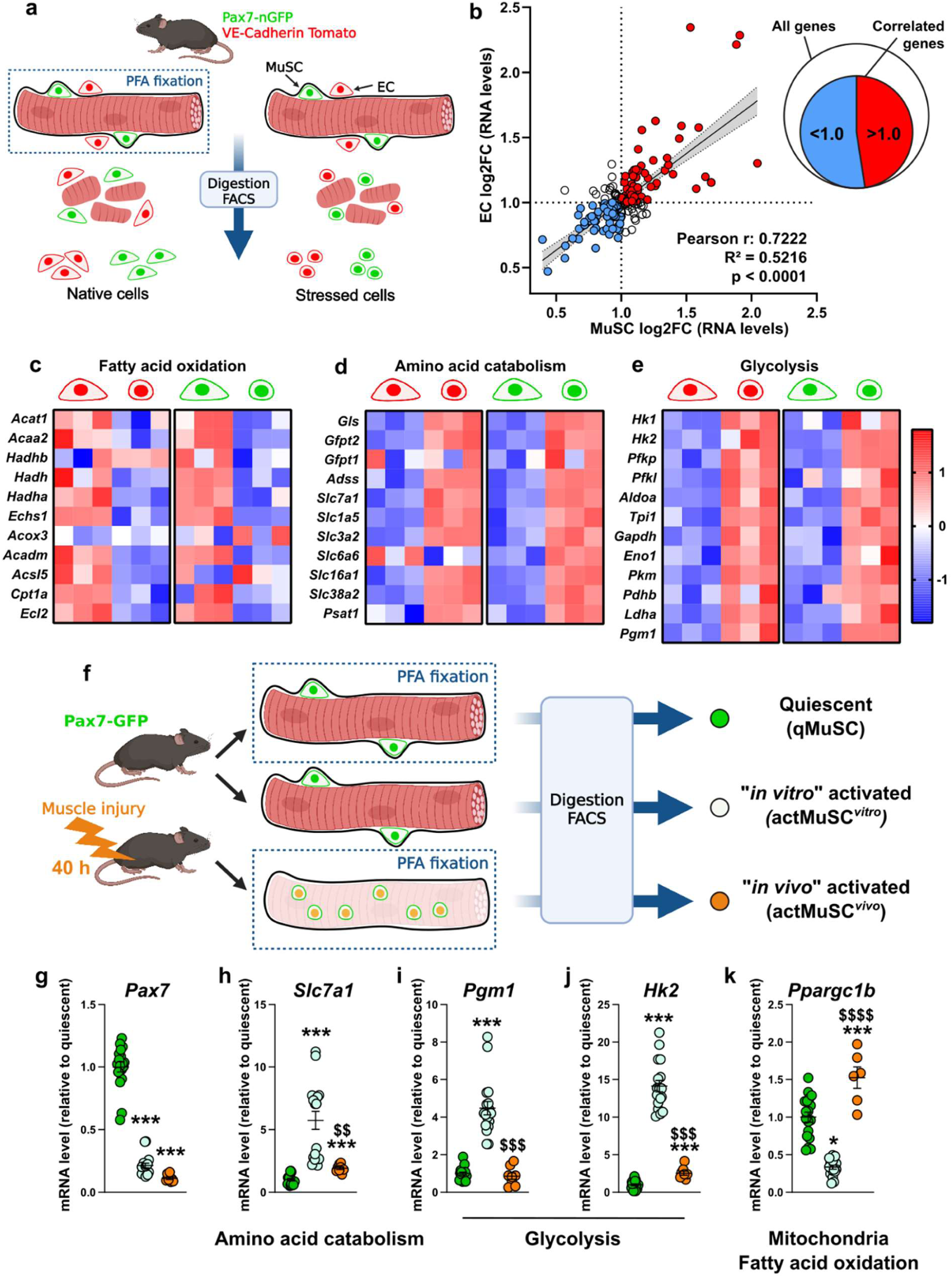
MuSC isolation alters their native metabolism. **a**, Experimental scheme for stressed or native MuSCs (nGFP^+^) and ECs (tomato^+^) isolated by FACS from fixed or unfixed tissue, respectively. **b,** Correlation of isolation-induced log2 fold change in metabolic gene expression between MuSCs and ECs. **c-e,** Heatmap showing the expression of genes involved in fatty acid oxidation (**c**), amino acid catabolism (**d**) or glycolysis (**e**) commonly altered during isolation in MuSCs and ECs. **f,** Experimental scheme for isolating MuSCs (nGFP^+^) by FACS from control or injured muscle, with or without prior fixation. **g-k,** mRNA expression levels of *Pax7* (**g**), *Slc7a1* (**h**), *Pgm1* (**i**), *Hk2* (**j**), and *Ppargc1b* (**k**), normalized to the geometric mean of three independent reference genes in MuSCs. Data are presented as mean ± SEM. *: *p* < 0.05; ***: *p* < 0.001 *vs* Quiescent. $$: *p* < 0.01; $$$: *p* < 0.001; $$$$: *p* < 0.0001 *vs* “*in vitro*” activated.

To evaluate if the changes in metabolic gene expression during MuSC isolation reflect their activation, we compared quiescent MuSCs (no injury, fixed before isolation, qMuSC) to MuSCs activated due to the isolation process (not fixed before isolation, thereafter called *“in vitro”* activation, actMuSC*^vitro^*) to MuSCs activated in the context of muscle injury (“*in vivo*” activated, actMuSC*^vivo^* fixed before their isolation, Fig. 1f). actMuSC*^vivo^* cells were collected 40 hours after injury, a time point selected to ensure a reduction in *Pax7* expression comparable to that of actMuSC*^vitro^* cells (Fig. 1g). The expression of *Slc7a1*, a transporter of amino acids, increased in both activated groups but to a significantly higher extent in actMuSC*^vitro^* cells (Fig. 1h). The expression of two genes involved in glycolysis, namely *Pgm1* and *Hk2*, strongly increased in actMuSC*^vitro^* cells, but not in actMuSC*^vivo^* cells for *Pgm1* or to a significantly lesser extent for *Hk2* (Fig. 1i-j). Finally, the expression of *Ppargc1b*, a regulator of fatty acid oxidation and mitochondrial respiration, was downregulated in actMuSC*^vitro^* cells and upregulated in actMuSC*^vivo^* cells (Fig. 1k). To further characterize the metabolic changes associated with MuSC activation, we performed metabolomic analyses on isolated MuSCs from control and injured muscles, seven days post-injury (Fig. S1b). To preserve a broad range of metabolites, MuSCs were not fixed, which prevented us from distinguishing between purely “*in vivo”* and “*in vitro”* activation. As a result, our analysis compared “*in vitro”* activated MuSCs to MuSCs exposed to both “*in vivo”* and “*in vitro”* activation. We observed a significant increase in numerous amino acids and metabolites derived from amino acid catabolism in activated MuSCs (Fig. S1c). However, expected changes in metabolites related to energy metabolism or fatty acid oxidation were not detected, suggesting that conventional metabolomics may overlook dynamic changes during MuSC activation or that such differences are lost during isolation. These results point out a methodological limitation of enzymatic dissociation technics in evaluating MuSC metabolism in a physiological context.

To avoid alteration of the MuSC microenvironment prior to the characterization of their metabolic profile, we optimized *in situ* staining of metabolic enzyme activity for muscle cryosections (Fig. 2a) (*23*, *24*). Combined with immunostaining of MuSC markers, this allows for quantitative metabolic profiling of MuSCs at the single-cell scale within their native microenvironment. We optimized or developed *in situ* enzymatic activity staining for 6 dehydrogenases catalyzing key reactions using specific substrates representative of 6 major metabolic pathways: 1. glucose-6-phosphate dehydrogenase (G6PD), a rate-limiting factor for the pentose phosphate pathway (PPP) (*25*); 2. glycerol-3-phosphate dehydrogenase (GPDH), a key producer of dihydroxyacetone phosphate (DHAP) to sustain glycolysis (*26*); 3. “oxidative” lactate dehydrogenase (LDH), whose activity constitutes the main source of pyruvate entering the tricarboxylic acid (TCA) cycle (*27*); 4. hydroxyacyl-CoA dehydrogenase/3-ketoacyl-CoA thiolase/enoyl-CoA hydratase subunit A (HADHA), a key enzyme for the β-oxidation of fatty acids (*28*); 5. glutamate dehydrogenase (GDH), involved in amino acid catabolism (*29*); and 6. succinate dehydrogenase (SDH), a mitochondrial enzyme involved in both the TCA cycle and the electron transport chain (Rutter, Mitochondrion 2010); (Fig. 2b-c). These assays allow for the quantification of the production of NAD(P)H/H+ or FADH2, the co-factor products of these dehydrogenases, which reduce light yellow nitroblue tetrazolium chloride (NBT) solution into violet insoluble formazan crystals, which bind to surrounding proteins and thereby allow precise localization of the investigated enzymatic activity at the cellular level. We validated the specificity of these reactions using validated pharmacological inhibitors of each enzyme-specific substrate (namely DHEA to inhibit G6PD, DHAP to inhibit GPDH, oxamate to inhibit LDH, trimetazidine to inhibit HADHA, EGCG to inhibit GDH and malonate to inhibit SDH) (Fig. 2c). Substrate specificity was further validated by applying increasing concentrations of substrates allowing us to quantify the dose-dependent signal intensity of enzymatic staining for each substrate in the entire tissue section (Fig. 2d). The resulting quantification curves followed a Michaelis-Menten equation, reflecting the catalytic reaction kinetics of each enzyme with a single substrate.

**Figure 2.**
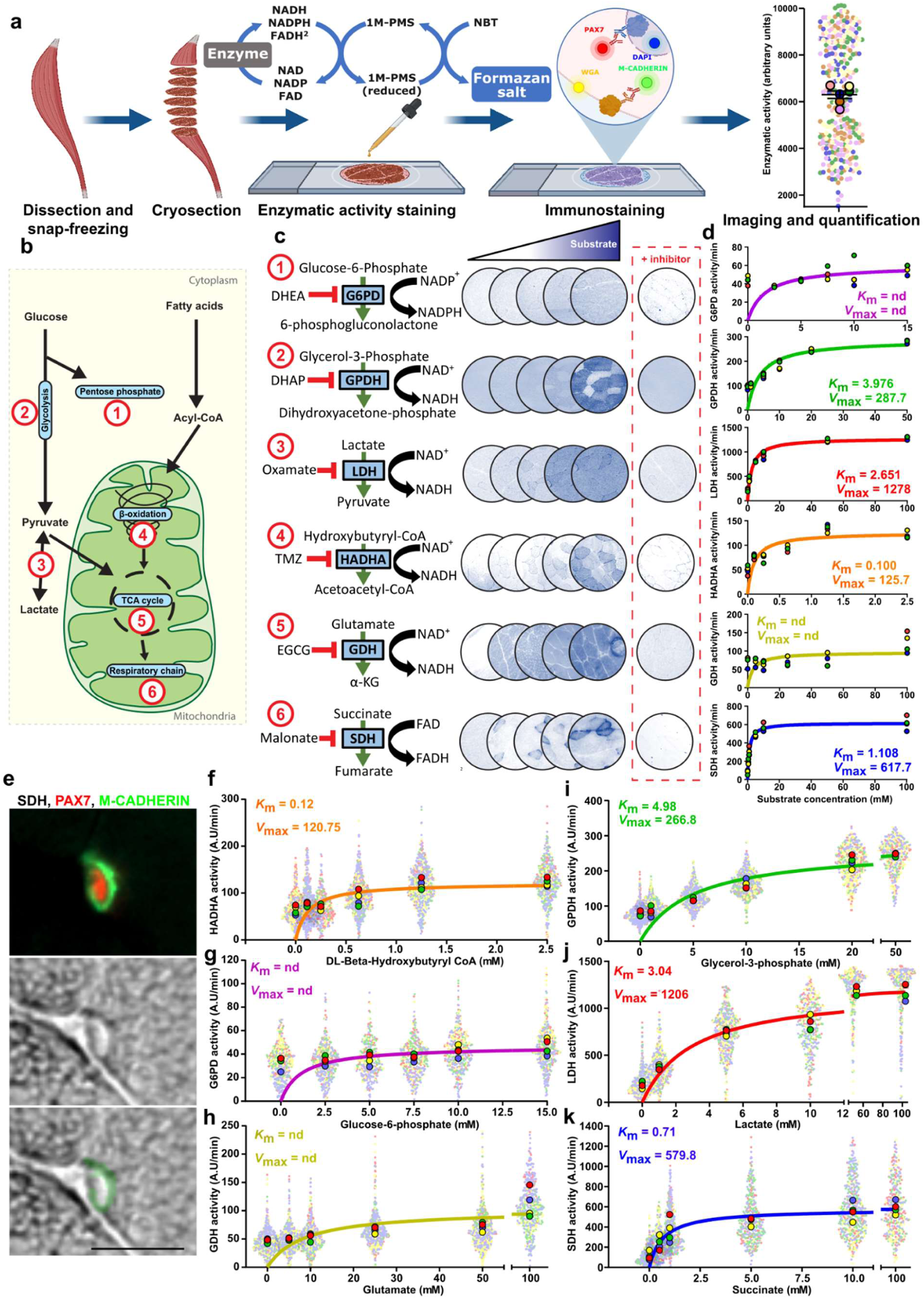
*In situ* enzymatic staining for metabolic profiling of MuSCs. **a**, Workflow for *in situ* enzymatic staining combined with immunostaining of MuSCs and myofibers. **b,** Schematic representation of metabolic pathways with the six selected enzymes. **c,** Scheme of enzymatic reactions and representative pictures of dose-response staining, with or without inhibitors. **d,** Michaelis-Menten kinetics of G6PD, GPDH, LDH, HADHA, GDH, and SDH, calculated based on whole-section quantification of dose-dependent activity staining. **e,** Compatibility of enzymatic staining with PAX7 and M-CADHERIN immunostaining. The middle panel shows a MuSC displaying low SDH staining in the nuclear region, surrounded by cytosolic regions with high SDH signal. The lower panel shows the mask used for quantification of MuSC enzymatic activity. Scale bar: 20 µm. **f-k,** Michaelis-Menten kinetics in MuSCs for HADHA (**f**), GPDH (**g**), G6PD (**h**), LDH (**i**), GDH (**j**), and SDH (**k**), based on quantification of dose-dependent activity staining. Each small, light dot represents a single MuSC or fiber, while each large, dark dot represents the mean for an individual mouse.

This approach enabled the determination of both the maximal enzymatic activity (*V*_max_) and the substrate affinity (*K*_m_), thereby confirming that each enzymatic activity can be quantitatively measured (Fig. 2d). Interestingly, despite extensive optimization of the experimental conditions, GDH and G6PD staining remained faint and yielded poor Michaelis–Menten fitting, indicating low enzymatic activity in resting TA muscle. Next, we tested the compatibility of *in situ* enzymatic activity staining (using SDH staining) with co-immunostainings using the following markers: PAX7 for MuSC nuclear staining, M-CADHERIN for MuSC plasma membrane staining, wheat germ agglutinin (WGA) for extracellular matrix counterstaining, and DAPI for nuclei staining, on cryosections from resting tibialis anterior (TA) (Fig. 2a, e). This co-staining enabled the quantification of multiple parameters within the same cryosections, including myofiber caliber and MuSC size alongside enzymatic activity. We then used this approach to specifically quantify enzymatic activity in MuSCs (Fig. 2f-k). Mean values were fitted to a Michaelis-Menten function to determine *V*_max_ and *K*_m_ of each enzyme. We obtained only faint staining for GDH and G6PD and low staining for HADHA, suggesting that the activity of these enzymes is low in quiescent MuSCs from resting TA muscle (Fig. 2f-h). Conversely, we obtained strong staining for SDH, GPDH, and LDH, revealing that these enzymes are highly active in MuSCs (Fig. 2i-k). Enzymatic properties measured across whole muscle sections or specifically within MuSCs were largely comparable, as reflected by similar *V*_max_ and *K*_m_ values (Fig. S2g–n). A notable exception was observed for SDH activity, whose *K*_m_ was significantly lower in MuSCs, indicating a higher substrate affinity in this cell population (Fig. S2k). Together, these results confirmed that *in situ* metabolic staining is specific, quantifiable, and thus represents a valid tool to assess the metabolic profile of MuSCs in their native environment.

The experimental approach confirmed the metabolic heterogeneity of myofibers in resting TA muscles, revealing small fibers with high oxidative staining (SDH and LDH) and large fibers exhibiting elevated glycolytic activity (GPDH) (Fig. 3a-d) as previously described (*30*). In this line, SDH and LDH oxidative staining were high in slow-twitch soleus muscles and low in fast-twitch superficial part of gastrocnemius muscles (Fig. 3g-i). Strikingly, we observed an unexpected heterogeneity in MuSC enzymatic activities in resting TA muscles using the SDH and GPDH staining (Fig. 3a-d), identifying the presence of both oxidative and glycolytic MuSCs. To validate this unanticipated result, we performed co-immunostainings of M-CADHERIN to stain the plasma membrane of MuSCs and COX IV to stain mitochondria as a marker of oxidative metabolism on freshly-isolated myofibers, from pre-fixed limb muscles to avoid dissociation-induced metabolic shift. We confirmed the presence of both oxidative and glycolytic MuSCs with dense or sparse mitochondrial networks, respectively, along two adjacent isolated myofibers (Fig. S3a). To identify which parameter best explains SDH staining heterogeneity, we performed a correlative analysis between SDH staining in MuSCs and all parameters measured by *in situ* metabolic profiling. We found that myofiber perimeter or area is inversely correlated with SDH staining, which is expected since oxidative fibers are known to be smaller (Fig. S3b). Remarkably, we noticed a strong and highly significant correlation between the intrinsic enzymatic activity of MuSCs and the enzymatic activity of neighboring myofiber in TA muscles (Fig. 3e-f, S3b). This implies that oxidative myofibers are surrounded by oxidative MuSCs, while glycolytic myofibers are surrounded by glycolytic MuSCs. To confirm these results, we performed SDH and GPDH staining on muscles with a more homogeneous metabolism, *i.e* the oxidative *soleus* and the glycolytic superficial *gastrocnemius*. Under these conditions, the correlation between myofiber and MuSC metabolic staining was preserved, indicating that the *soleus*, which contains a high proportion of oxidative fibers also contains a high proportion in oxidative MuSCs, and that the superficial *gastrocnemius*, which contains high proportions of glycolytic fibers, also contains a high proportion of glycolytic MuSCs (Fig. 3g-i and S3c-d). For both myofiber and MuSC oxidative activities, analysis of variance demonstrated a significant difference between the TA and the *soleus* as well as between the TA and the superficial *gastrocnemius*, further confirming that the heterogeneity of MuSC metabolism is linked to myofiber metabolic heterogeneity (Fig. 3g-i).

**Figure 3.**
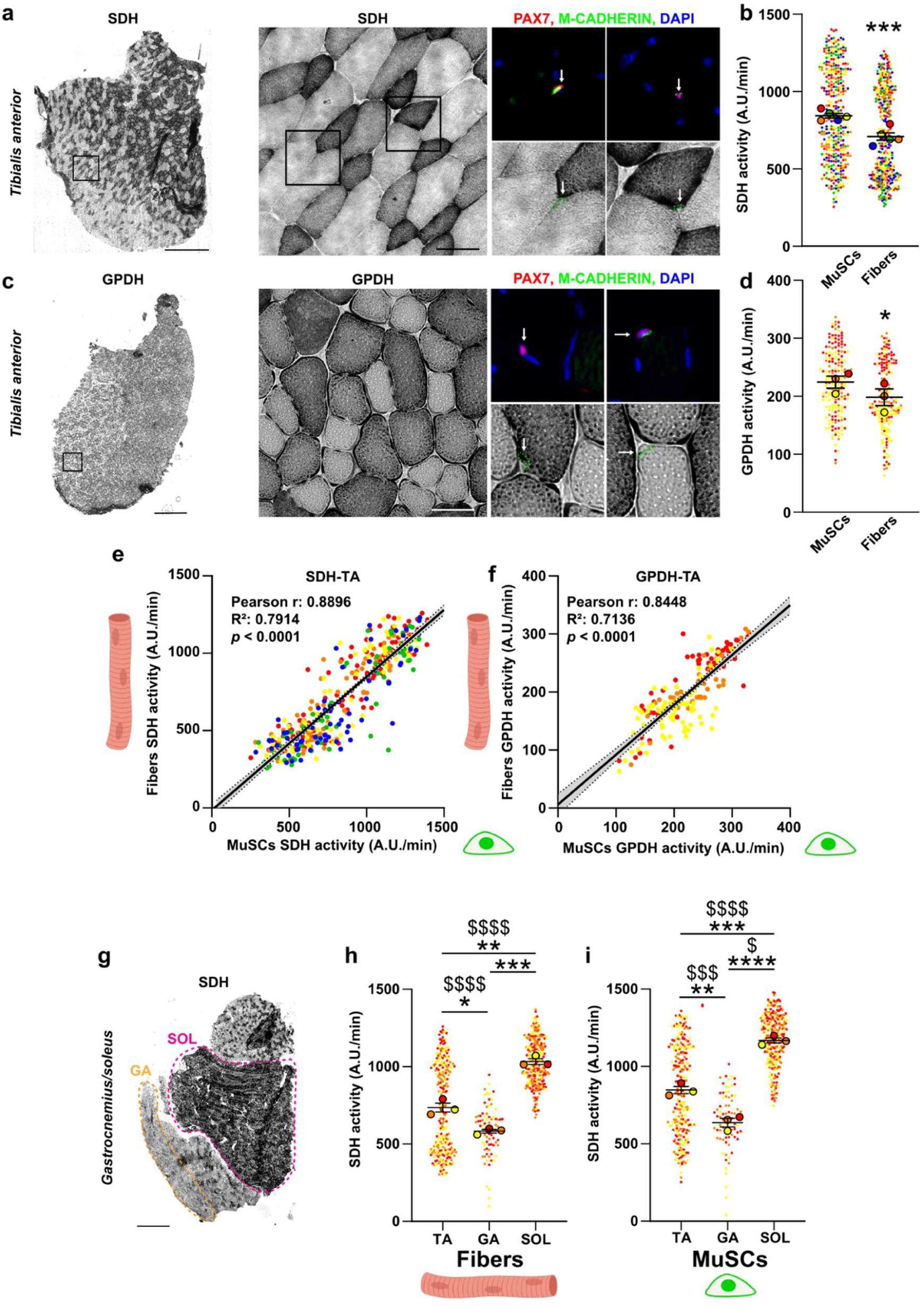
ISE reveals MuSC metabolic heterogeneity and the correlation between MuSC and myofiber metabolism. **a**, Representative SDH staining of an entire tibialis anterior section (left) and a higher magnification view (right), with cropped sections showing immunostaining (top) for M-cadherin (green), Pax7 (red), and DAPI (blue) or SDH staining (grey, bottom) with the quantified MuSC region highlighted in green. **b,** Quantification of SDH activity in MuSCs (left) and fibers (right). Each small, light dot represents a single MuSC or fiber, while each large, dark dot represents the mean for an individual mouse with error bars showing the SEM. The activity quantification reveals two distinct subpopulations of fibers and MuSCs. **c,** Representative GPDH staining of an entire tibialis anterior section (left) and a higher magnification view (right), with cropped sections showing immunostaining (top) for M-cadherin (green), Pax7 (red), and DAPI (blue) or GPDH staining (grey, bottom) with the quantified MuSC region highlighted in green. **d,** Quantification of GPDH activity in MuSCs (left) and fibers (right). Each small, light dot represents a single MuSC or fiber, while each large, dark dot represents the mean for an individual mouse with error bars showing the SEM. The activity quantification reveals two distinct subpopulations of fibers and MuSCs. **e-f,** Linear regression analysis of SDH (**e**) and GPDH (**f**) activity in *tibialis anterior* fibers (y-axis) and associated MuSCs (x-axis) with Pearson correlation scores, revealing a correlation between the metabolic profiles of fibers and associated MuSCs. **g,** Representative SDH staining of an entire *gastrocnemius* and *soleus* section. The superficial glycolytic region of the *gastrocnemius* (GA) is outlined in yellow, and the oxidative *soleus* (SOL) is outlined in pink. **h-i,** Quantification of SDH activity in fibers (**h**) and MuSCs (**i**) in the *tibialis anterior* (TA), superficial *gastrocnemius* (GA), and *soleus* (SOL). For b and d, *: *p* < 0.05; ***: *p* < 0.001 *vs* fibers. For h and i, *: *p* < 0.05; **: *p* < 0.01; ***: *p* < 0.001; ****: *p* < 0.0001 for differences of means (T-tests); $: *p* < 0.05; $$$: *p* < 0.001; $$$$: *p* < 0.0001 for differences of variances (F-tests). Scale bars: 500 µm on entire sections, 50 µm on higher magnifications.

To characterize more precisely the bidirectional interplay between myofiber and MuSC metabolic profiles, we used a surgical model of muscle denervation known to alter myofiber metabolism by reducing contractile activity (*31*) (Fig. 4a). We first validated the expected metabolic shift of denervated myofibers in TA towards a more oxidative phenotype 3 weeks post-denervation, using *in situ* SDH activity staining (Fig. 4b-c). Of interest, a similar oxidative metabolic shift was observed in MuSCs in denervated TAs (Fig. 4d-e), and the correlation between myofiber and MuSC enzymatic activities was preserved in denervated muscles (Fig. 4f). These findings support the hypothesis that myofiber metabolism controls MuSC metabolism *in vivo.* To further confirm that a change in myofiber metabolism induces a change in MuSC metabolism, we used *Tsc1*mKO mice, a model in which TSC1 is specifically invalidated in myofibers. In *Tsc1*mKO mice, the mTORC1 pathway is constantly activated and leads to increased oxidative capacity in TA myofibers (Fig. 4g) (*32*). In this model, we confirmed the increase in SDH activity in myofibers from *Tsc1*mKO mice (Fig. 4h-i) and observed a similar metabolic shift in MuSC metabolism (Fig. 4j-k), with the correlation between myofiber and MuSC enzymatic activities being preserved under these conditions (Fig. 4l). Together, these results demonstrate a previously unknown role of myofibers in regulating MuSC metabolism.

**Figure 4.**
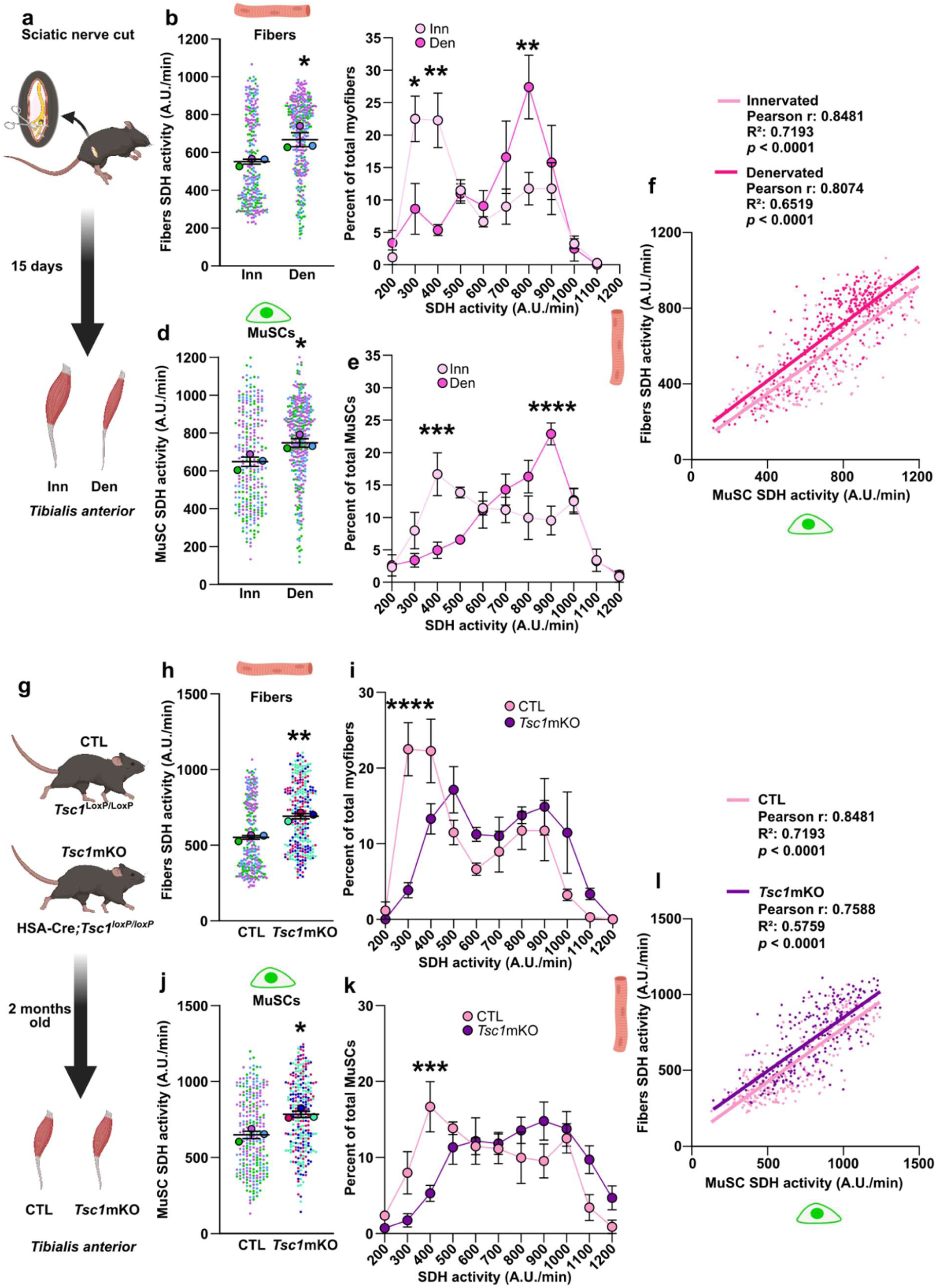
Metabolic shift in myofibers induces a similar metabolic shift in MuSCs. **a**, Workflow for muscle denervation**. b-e,** Quantification of SDH activity in fibers (**b-c**) and MuSCs (**d-e**) in denervated (Den, dark pink) or innervated contralateral tibialis anterior muscle (Inn, light pink). Each small, light dot represents a single MuSC or fiber, while each large, dark dot represents the mean for an individual mouse with error bars showing the SEM. **c** and **e** show the distribution of cells (as % of the total population), according to their SDH activity. **f,** Linear regression analysis of SDH activity in innervated (light pink) and denervated (dark pink) tibialis anterior fibers (y-axis) and associated MuSCs (x-axis) with Pearson correlation scores, revealing a correlation between the metabolic profiles of fibers and associated MuSCs in both conditions. **g,** Workflow for myofiber specific *Tsc1* deletion**. h-k,** Quantification of SDH activity in fibers (**h-i**) and MuSCs (**j-k**) in tibialis anterior muscle from control (CTL) or *Tsc1*mKO. Each small, light dot represents a single MuSC or fiber, while each large, dark dot represents the mean for an individual mouse with error bars showing the SEM. **i** and **k** show the distribution of cells (as % of the total population), according to their SDH activity. **l,** Linear regression analysis of SDH activity in control (light pink) and *Tsc1*mKO (purple) tibialis anterior fibers (y-axis) and associated MuSCs (x-axis) with Pearson correlation scores, revealing a correlation between the metabolic profiles of fibers and associated MuSCs in both conditions.

Finally, we investigated the kinetics of myofiber-dependent regulation of MuSC metabolism during muscle regeneration. To this end, we mapped SDH activity at 5, 7, 14, and 28 days after injury focusing on PAX7^+^ cells and neighboring regenerating fibers in regenerating regions characterized by the presence of centronucleated myofibers. (Fig. 5a). We noticed a drop in SDH activity in regenerating fibers at day 5 and 7 post-injury compared to intact fibers (Fig. 5b). The resulting disruption of MuSC metabolic niche results in a concomitant decrease of SDH activity in PAX7^+^ cells at days 5 and 7 post-injury (Fig. 5c). By day 14, SDH activity had returned to baseline in both cell types. Interestingly, SDH activity was even significantly elevated in both MuSCs and fully regenerated fibers at day 28 post-injury, compared with day 0 (Fig. 5b, c). We next evaluated if the metabolic interplay between the myofiber and the adjacent MuSC was also found in the context of muscle regeneration. We observed a strong correlation between the metabolic state of myofibers and that of associated PAX7^+^ cells across all analyzed time points (Fig. S5a-e). In addition to PAX7⁺ cells, we identified a population of PAX7⁻/M-cadherin⁺ mononucleated cells that were not associated to myofibers, and were classified as differentiated cells, at days 5, 7, and 14 post-injury. In 2 mice out of 5, differentiated cells were still detectable at day 14 post-injury (Fig.5d). Quantification revealed that in contrast to PAX7^+^ cells in which SDH activity is increasing, differentiated cells consistently displayed low SDH activity at days 5, 7, and 14 (Fig. 5e). Consequently, we found that SDH activity in myofibers was not significantly correlated to that of differentiated cells (Fig. 5f). These results suggest that direct cell-cell contact is required to synchronize MuSC and myofiber metabolism. Finally, analysis of cell distribution revealed that MuSCs and myofibers display heterogeneous SDH activity, whereas differentiated cells exhibited uniformly low SDH activity (Fig. S5f).

**Figure 5.**
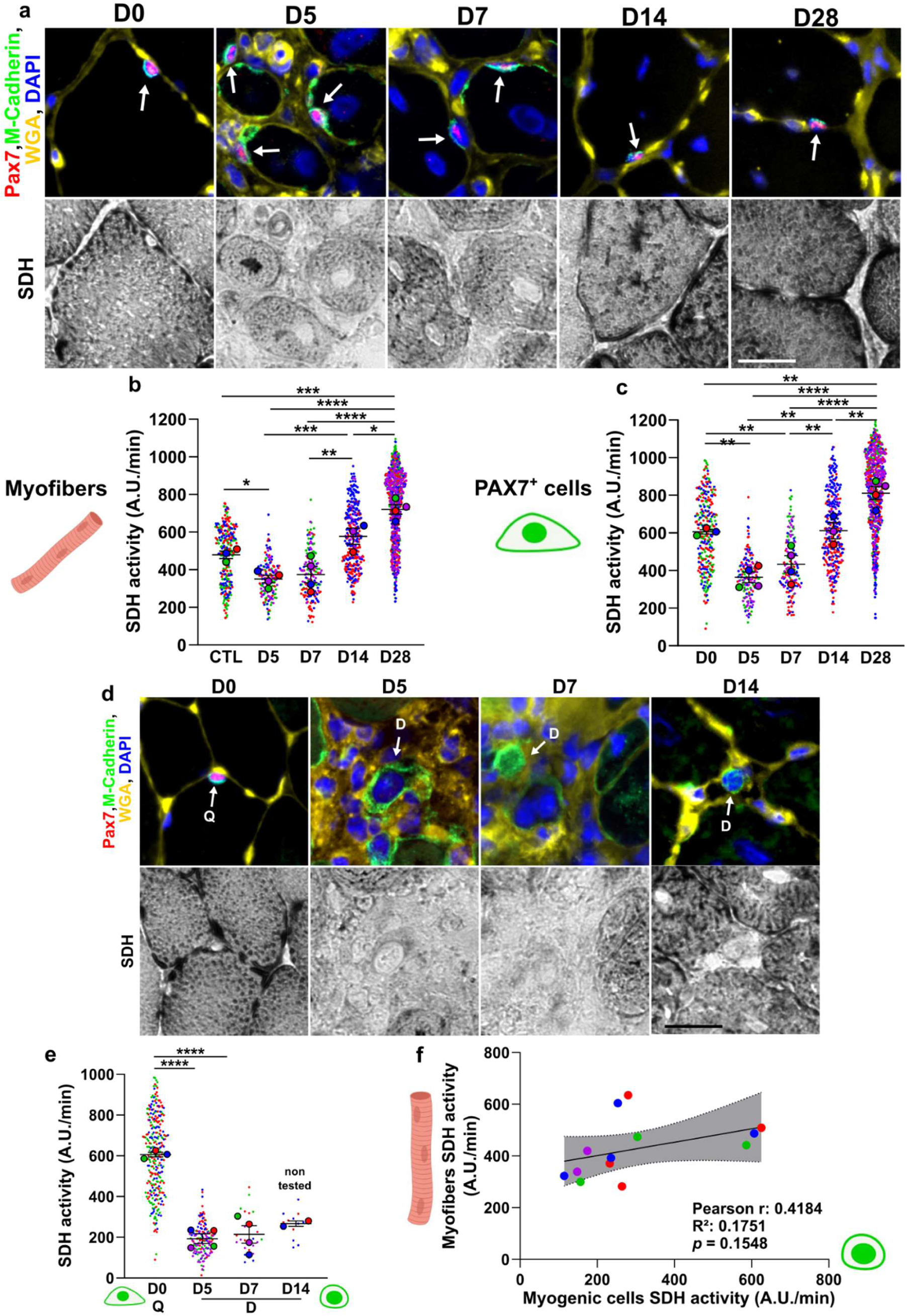
Mapping of oxidative metabolism during muscle regeneration. **a**, Representative views displaying immunostaining (top) for M-cadherin (green), PAX7 (red), WGA (yellow), and DAPI (blue), alongside SDH enzymatic staining (grey, bottom) throughout the process of muscle regeneration. Arrows indicate PAX7^+^ cells. Scale bar: 30 µm. **b-c,** Quantification of mean SDH enzymatic staining in myofibers (**b**) and associated PAX7^+^ cells (**c**), during the process of muscle regeneration. Each small, light dot represents a single cell, while each large, dark dot represents the mean for an individual mouse with error bars showing the SEM. **d**, Representative images showing immunostaining (top) for M-cadherin (green), PAX7 (red), WGA (yellow), and DAPI (blue), alongside SDH enzymatic staining (grey, bottom) at D0 (left) and D5 after injury (right). Arrows indicate quiescent (Q) and differentiated (D) cells. **e**, Quantification of mean SDH activity in Q cells (at D0) and D cells (at D5, D7, and D14). Each small, light dot represents a single cell, while each large, dark dot represents the mean for an individual mouse, with error bars showing the SEM. **f,** Linear regression analyses of average SDH activity in fibers (y-axis) and D cells (x-axis) in *tibialis anterior*, with Pearson correlation scores. Scale bars: 20 µm.

## Discussion

It is well-established that the primary regulator of cellular metabolism is the availability of substrates and co-substrates within the surrounding microenvironment (*1–4*, *33*). However, this crucial parameter is often overlooked due to significant technical limitations. In the field of stem cell research, early evidence regarding the impact of the microenvironment on metabolism was obtained from studies on MuSCs. Four independent laboratories developed methods to identify the epigenetic and transcriptomic changes that occur during the isolation of MuSCs (*34–37*). These four studies revealed a set of genes associated with MuSC activation, proliferation, and exit from quiescence. Additionally, they identified numerous genes involved in metabolic pathways, along with stress-related genes, which were likely upregulated as a consequence of the isolation process. Importantly, similar gene expression changes were observed during cell isolation from other tissues, emphasizing that studying cell function within its native microenvironment remains a key, yet often neglected, consideration (*22*). These findings further highlighted the presence of artefacts in large-scale datasets, including single-cell RNA-seq resources, emphasizing the need for caution when interpreting such data.

In this study, we demonstrated that cell isolation also distorts the native metabolic state of cells, independently of the associated changes in cell fate. Our observations suggest that altered expression of metabolic genes reflects the composition of the isolation buffer in terms of nutrients. Similar observations have been reported in cancer research, where glutamine is a predominant substrate fueling the TCA cycle in cultured cancer cells, but its consumption appears to be minimal in tumors *in vivo* (*38*). This finding opens new opportunities to develop media formulations based on nutrients composition that selectively activate specific metabolic pathways to influence cell fate. This could have particular relevance in the context of cell isolation for therapeutic applications. Furthermore, our results suggest the importance of using well-defined media in academic research and call for manufacturers to disclose the compositions of commercial media transparently. Instead, the development and sharing of standardized media formulations could significantly accelerate discoveries in the field of cell therapy by enabling more reproducible and biologically relevant studies.

Studying metabolic pathways activity is particularly challenging due to its highly dynamic nature. Major shifts in metabolic fluxes, such as a rapid switch from fatty acid oxidation to glycolysis, can occur within seconds to maintain energy supply, without any change in protein expression. Consequently, while transcriptomic or proteomic data can provide insights into long-term metabolic reprogramming, they are insufficient for accurately capturing rapid and transient metabolic changes. Furthermore, these omics approaches cannot quantify the actual enzymatic activity of a specific metabolic pathway or the metabolic capacity of a given cell. The most direct method for evaluating metabolic fluxes is fluxomics, which combines *in vivo* isotope labeling with metabolomics to track substrate fate. However, this technique requires cell isolation and fixation prior to isolation is not compatible with mass spectrometry-based analyses, as fixation interferes with detection of many metabolites. While spatially resolved fluxomics approaches are emerging, their resolution (typically ∼50 µm) remains insufficient for the analysis of very small flat cells like MuSCs (∼10 µm in diameter).

To overcome this, we developed a novel method for *in situ* metabolic profiling of MuSCs metabolism by staining for the activity of metabolic enzymes on muscle cryosections, combined with MuSC-specific immunostaining. Because rapid metabolic adjustments rely on modulation of enzymatic activity, particularly through allosteric regulation (*1–3*), enzymatic activity has long been considered a reliable indicator of metabolic capacity in muscle research (*39*). Moreover, enzymatic activities can reflect not just the function of individual enzymes, but also the coordination between them. In our experimental setup, we found that inhibiting Complex IV affected Complex II activity, highlighting how this method can provide insight into coupled enzymatic fluxes. Thus, such staining provides the advantage of monitoring MuSC metabolism in their native environment at the single cell scale and is based on a readout that is more relevant than transcriptomic or proteomic data. Similar approaches have been successfully developed to characterize the metabolic configuration of immune cells in tumors (*23*).

One unexpected observation from the *in situ* enzymatic activity assays was the marked metabolic heterogeneity among quiescent MuSCs. Within this heterogeneous population, we identified a substantial subpopulation, comprising approximately 50% of the total MuSCs, with notably high metabolic activity. As these cells were analyzed under healthy, non-injured conditions, it is unlikely that they correspond to activated or self-renewing MuSCs; rather, they appear to be metabolically active quiescent cells. This finding suggests that the maintenance of quiescence is not necessarily a passive, low-energy state but may involve active processes that impose a significant energetic cost on the cell. While this hypothesis requires further validation and the underlying mechanisms remain to be elucidated, it opens new perspectives for understanding MuSC function, and dysfunction, particularly in pathological contexts characterized by MuSC pool exhaustion. Based on these observations, we propose that the metabolic heterogeneity observed in quiescent MuSCs reflects variability in the depth of quiescence. Several studies have suggested a spectrum of quiescent states, ranging from a shallow state, characterized by cells primed for rapid activation, to a deep quiescent state, where cells are highly resistant to activating stimuli (*40–43*). In MuSC this gradation has been linked to differential Pax7 expression levels and associated metabolic profiles (*15*, *44*, *45*). Rodgers and colleagues also described a shallow quiescent state, termed “G_Alert,” marked by specific metabolic changes (*46*). Notably, the depth of quiescence has been linked to lysosomal function, which depends on ATP consumption for acidification, reinforcing the idea that maintaining quiescence may involve significant metabolic investment (*43*).

Although metabolism clearly plays a role in modulating MuSC states, the precise relationship between specific metabolic programs and the depth of quiescence remains poorly defined. For instance, it is not yet clear whether a more oxidative or glycolytic profile actively promotes a shallow or deep quiescent state, or whether these metabolic traits are consequences rather than drivers of cell state. This uncertainty highlights the need for further studies dissecting causality between metabolic activity and cell fate. We propose that the metabolic heterogeneity within quiescent MuSCs is essential to support this spectrum of quiescent states. Such diversity would serve a dual purpose: maintaining a long-term, self-renewing MuSC pool while preserving a subset of cells primed for rapid activation. Interestingly, metabolic heterogeneity among MuSCs is markedly reduced after injury, with nearly all cells adopting a similar metabolic profile. Unexpectedly, this homogeneity persists for an extended period, as regenerated muscles at 28 days post-injury still display uniformly high oxidative capacities in both myofibers and associated MuSCs. We propose that this sustained oxidative state reflects the delayed maturation of glycolytic, fast-twitch fibers, which require more time to reestablish (*47*, *48*). In this context, it is particularly interesting to consider the correlation between myofiber metabolism and the metabolic profile of their associated MuSCs. This relationship could influence the regenerative or growth potential of different muscles. Supporting this idea, Collins and colleagues reported higher grafting efficiency for MuSCs isolated from the oxidative *soleus* muscle compared to those from the more glycolytic EDL muscle (*49*). Since quiescence depth is associated with grafting success, these findings suggest that MuSCs from oxidative muscles may be more deeply quiescent. Consistent with this, recent work has shown that the proportions of PAX7 High and PAX7 Low MuSCs differ between oxidative and glycolytic muscles (*15*). A metabolic role for myofibers could also be involved in the findings that dysfunctional MuSCs in aged or dystrophic mice models recover their function when grafted into young healthy hosts (Boldrin, Zammit, et Morgan 2015; Novak et al. 2021).

Finally, although the molecular mechanisms underlying myofiber regulation of MuSC metabolism remain to be elucidated, our study provides some insights. We identified that metabolic state of myogenic cells correlates with myofiber metabolism only after establishing contact. This contrasts with cells lacking such contact, in which metabolic status was determined primarily by their activation state. We speculate that physical interaction with myofibers may either activate membrane receptors or facilitate metabolite transfer, thereby reprogramming MuSC metabolism. Further studies will be needed to answer this question.

In conclusion, *in situ* enzymatic staining represents a powerful and accessible tool to uncover novel aspects of metabolic regulation. Its compatibility with frozen samples makes it particularly suitable for the analysis of human biopsies, offering strong translational potential. We anticipate that this method can be readily adapted to a wide range of cell types and tissues, greatly expanding its utility across biomedical research. Beyond its potential as a diagnostic or therapeutic tool, this approach opens exciting avenues for fundamental discoveries. Applied to MuSCs, our method reveals the critical role of the local microenvironment in shaping MuSC metabolism. We uncover an unexpected degree of metabolic heterogeneity among quiescent MuSCs and demonstrate that their metabolic state is directly influenced by neighboring myofibers. These findings highlight metabolism not merely as a passive reflection of cell state, but as an actively regulated property shaped by cell–cell interactions, adding a new dimension to our understanding of stem cell biology.

## Methods

### Animals

Mouse lines used in this study were described previously: *Tg:Pax7-nGFP* mice (*50*), *Tg:Cdh5-CreERT2/+* mice (*51*) R26^(stop)Tomato/+^ mice (*52*). To visualize PAX7^+^ cells with nGFP and VE-CADHERIN^+^ endothelial cells (ECs) with Tomato, *Tg:Cdh5-CreERT2/+* mice were intercrossed with R26^(stop)Tomato/+^ mice. Then, *Tg:Cdh5-CreERT2/+; R26*^(stop)Tomato/+^ generated mice were intercrossed with *Tg:Pax7-nGFP* mice in order to obtain *Tg:Pax7-nGFP;Cdh5-CreERT2/+;R26*^(stop)Tomato/+^ mice and to visualize both nGFP^+^ MuSCs and Tomato^+^ ECs. To induce recombination, 6-week-old *Tg:Pax7-nGFP/+*; *Tg:Cdh5-CreERT2/+*; *R26*^fl(stop)fl Tomato/+^ mice were maintained on a low-phytoestrogen tamoxifen diet (Altromin 1324P, Genetsil) for 10 consecutive days. Eight- to 12-week-old mice were used to isolate MuSCs and ECs. Generation and genotyping of TSCmKO transgenic mice were described previously (*31*). Control mice for TSCmKO mice were littermates that were floxed for *Tsc1* but did not express Cre-recombinase. For muscle denervation, sciatic nerve was cut unilaterally on mice anesthetized with isoflurane, as previously described (*32*). The contralateral leg serves as control. For muscle injury, mice were anaesthetized with isoflurane. One TA muscle was injected with 50μl of the cobra venom cardiotoxin (CTX) (10μM, Latoxan laboratory; # L8102). All mouse experiments were conducted in accordance with the European Union guidelines on animal experimentation and approved by the Veterinary Office of the Canton of Geneva (application number GE220/GE227) and the ethics committee at the relevant French Ministries (Project No: 16-062) as appropriate. Quickly after euthanasia, muscles from mice were dissected and snap-frozen in isopentane cooled in liquid nitrogen and then stored at −80°C. eight-micrometer-thick transverse sections were used for ISE.

### *In Situ* Enzymatic staining (ISE)

Eight-micrometer-thick transverse sections were used for ISE. Immediately before performing the experiment, tissue sections were thawed for two minutes at room temperature. Pre-heated (37°C) assay medium containing the enzyme-specific substrate and coenzymes was applied to cover the whole tissue section. Enzyme reactions were carried out at 37°C, protected from light for 8 min (SDH, LDH) or 30 min (GPDH, G6PD, GDH; GAPDH, HADHA). Specific buffers were prepared as follows: For the SDH, GPDH, LDH, G6PD and GDH assays the buffer was 0.1 M Tris-maleate buffer pH 7.5, for the detection of GAPDH activity it was 0.1 M Tris-maleate buffer pH 8.0 and for the HADHA activity assay it was 0.1 M pyrophosphate buffer pH 7.3. In these enzyme specific buffers, ten percent polyvinyl alcohol was dissolved by stirring in a water bath at 60°C until the mixture was clear. Assay media were freshly prepared and contained for SDH, GAPDH, LDH, G6PD and GDH activity assays 0.45 mM 1-methoxyphenanzine methosulfate, 5 mM sodium azide and 4.95 mM nitroblue tetrazolium chloride (pre-dissolved in 70% dimethylformamide). The assay media for GPDH activities contained 1 mM menadione instead of 1-methoxyphenanzine methosulfate. The assay media also contained 15 mM G6P, 0.8 mM NADP+ and 4 mM MgCl_2_ for the analysis of G6PD activity; 2.5 mM glyceraldehyde-3-phosphate (pre-dissolved in 0.01M HCl) 2.5 mM of dihydroxyacetone phosphate and 3 mM NAD+ for the analysis of GAPDH activity; 10 mM sodium lactate and 2.5 mM NAD+ for the analysis of LDH activity; 50 mM sodium succinate for the analysis of SDH activity; 100 mM glutamate, 2.5 mM NAD+ for the analysis of GDH activity; 50 mM of Rac-G1P for the analysis of GPDH activity; 2.5 mM DL-β-hydroxybutyryl CoA, 2.5 mM NAD+ for the analysis of HADHA activity. Negative control reactions were performed in absence of substrate or presence of 80 mM DHEA (for G6PD inhibition), 40 mM sodium iodoacetate (for GAPDH inhibition), 200 mM sodium oxamate (for LDH inhibition), 880 µM of trimetazidine (for HADHA inhibition) 50 mM of dihydroxyacetone phosphate (for GPDH inhibition) 50 µM of epigallocatechin-3-gallate (for GDH inhibition) or 250 mM malonic acid (for SDH inhibition). Enzyme inhibitors were applied at very high concentrations for maximum inhibition, irrespective of enzyme selectivity concerns as the reaction specificity of the activity assays are defined by the provided substrates.

### Immunostaining after ISE

Immediately after ISE, sections were washed with ice-cold PBS and fixed with 4% PFA for 8 min at RT. After a wash with PBS, PFA was quenched with glycine (0.1M pH7.4 in PBS) during 8 min at RT and sections are subsequently incubated for 8 min in cold (−20°C) methanol. After a wash with PBS sections underwent antigen retrieval for 7 min in boiling citric acid buffer (0.01M, pH6). Sections were then incubated in blocking solution for one hour at RT (BSA 3% (IgG free – Jackson Immunoresearch), 0.5% triton and 1:100 AffiniPure Fragment Mouse Fab (Jackson Immunoresearch) in PBS). Primary antibodies were then incubated overnight, followed by three washes, and incubation with the corresponding secondary antibodies (Jackson ImmunoResearch), together with wheat – germ agglutinin-AlexaFluor647 (Invitrogen). For PAX7 detection, a biotinylated antibody was used followed by fluorescent streptavidin (Jackson Immunoresearch) incubation to amplify the signal. Sections were then washed and mounted in Vectashield DAPI (Vector). Images were recorded using an Axio Imager M2 with ×20 objective (Zeiss).

### Immunostaining of single isolated fibers

*Tibialis anterior* muscles were removed, and the entire leg fixed in 2% PFA for 25 min. The leg was washed, and EDL bundles were cut, and single fibers isolated with forceps under binocular microscope from PFA-fixed EDL muscle. Single fibers were then permeabilized with PBS, 3% BSA, 5% Triton and AffiniPure Mouse IgG Fab Fragments (1:100, Jackson ImmunoResearch). Primary antibodies were incubated overnight, followed by over day washing, and incubation with the corresponding secondary antibodies (Jackson ImmunoResearch), together with wheat –germ agglutinin-AlexaFluor647 (Invitrogen) at RT. Fibers were then washed overnight and mounted in Vectashield DAPI (Vector). Images were recorded using an LSM-800 confocal microscope with ×63 objective (Zeiss).

### Antibodies

The following primary antibodies were used: M-Cadherin (Cell Signaling Technology, #240491, 1:250 for IF); PAX7 (Development Studies Hybridoma Bank, AB_528428, 1:50 for IF); COXIV (Abcam, ab14744, 1/250 for IF).

### MuSC and ECs isolation

“Quiescent” and “*in vivo* activated” samples were prepared according to the original *iSiFi* protocol (*37*). Briefly, the samples were fixed for 1 h in 0.5% PFA, washed 3 times in ice-cold PBS and digested for 2 h at 37°C under shaking in 2X digestion solution: Dispase II, 6 U/ml (Roche, 4942078001), Collagenase A 1 U/ml (Roche, 10103586001) and 0.2% BSA:DMEM. Stressed cells or “*In vitro*” activated samples were digested for 120 min in 1X digestion solution: Dispase II, 3 U/ml (Roche, 4942078001), Collagenase A 0.5 U/ml (Roche, 10103586001) and 0.2% BSA/DMEM at 37°C with agitation. The samples were then spun down (600xg, 5 min) and resuspended in ice-cold 0.5% PFA and incubated for 1 h at 4°C with gentle agitation to fix the cells. At this point, all the samples were filled with DMEM (GIBCO, 41966-029) and filtered successively through 100 μm and 70 μm cell strainers (MACS, 130-110-917 and 130-110-916 respectively) then span down (600xg, 5 min), resuspended in ice-cold DMEM and filtered through 40 μm cell strainer (Corning, 352340) before a last centrifugation (600xg, 5 min) and resuspension in 0.2% BSA/DMEM for cell sorting.

### RNA Extraction and qRT-PCR

RNA was extracted using the RecoverAll Kit for FFPE (Ambion, #AM1975) following the manufacturer’s guidelines, with slight modifications: the incubation step at 50°C was performed for 1 hr instead of 15 min to improve RNA yield, and the incubation step at 80°C was omitted, as it deteriorated the quality of the recovered RNA. Reverse transcription was performed using the SuperScript III Reverse Transcriptase kit (Thermo Fisher Scientific, #18080093) with random primers, following the manufacturer’s guidelines. qPCR was performed using the Power SYBR Green PCR Master Mix (Applied Biosystems, #4367659). All reactions were run in triplicates and normalized to the expression *Rplpo* as housekeeping genes. The following primers were used: Pax7(for)GCGAGAAGAAAGCCAAACAC; Pax7(rev)CGGGTTCTGATTCCACATCT; Slc7a1(for) AAACCCCGGACATATTTGCT; Slc7a1(rev)ACCATGGCTGACTCCTTCAC; Pgm1(for) TCAGTGACCTGAAGCAGAGG; Pgm1(rev)CTGGGTGATAAATGCAGTCG; Hk2(for)GACCACATTGTCCAGTGCAT; HK2(rev)TTTGTCCACTTGAGGAGGATG; Ppargc1b(for)TGGAAAGCCCCTGTGAGAGT; ppargc1b(rev)TTGTATGGAGGTGTGGTGGG

### RNA-sequencing

RNA-Sequencing RNA quality and yield were assessed by the RNA integrity number (RIN) algorithm, using the 2100 Bioanalyzer. Directional libraries were prepared using the Smarter Stranded Total RNA-Seq kit-Pico Input Mammalian kit following the manufacturer’s instructions (Clontech, 635005). The quality of all libraries was verified with the DNA-1000 kit (Agilent) on a 2100 Bioanalyzer and quantification was performed with Quant-It assays on a Qubit 3.0 fluorometer (Invitrogen). Clusters were generated for the resulting libraries, with the Illumina HiSeq SR Cluster Kit v4 reagents. Sequencing was performed using the Illumina HiSeq 2500 system and HiSeq SBS kit v4 reagents. Runs were carried out over 65 cycles, including seven indexing cycles, to obtain 65-bp single-end reads. Sequencing data were then processed with the Illumina Pipeline software, Casava version 1.9. Reads were cleaned of adapter sequences and low-quality sequences using an in-house program (https://github.com/baj12/clean_ngs). Only sequences at least 25nt in length were considered for further analysis. STAR version 2.5.0a (*53*), with default parameters was used for alignment on the reference genome (GRCm38 from Ensembl database). Genes were counted using featureCounts version 1.4.6-p3 (*54*) from Subreads package (parameters: -t gene -s 0 -O). Count data were analyzed using R version 3.3.1(R Core Team, 2016) and the Bioconductor package DESeq2 version 1.12.3 (*55*). The normalization and the dispersion estimation were performed with DESeq2 using the default parameters, but statistical tests for differential expression were performed without applying the independent filtering algorithm.

### Broad-scale targeted metabolomics

The metabolomic analysis was performed by the Metabolomics and Lipidomics Platform of Lausanne’s University, as previously described (*56*). Briefly, broad-scale targeted metabolomic analysis was performed on FACS-sorted MuSCs cells isolated from control or CTX-injured TAs. Approximately 150,000 freshly isolated MuSCs per condition were collected in cold MeOH:H_2_O (9:1). Methanol extracts were dried, resuspended in 75 µl of MeOH:H_2_O (4:1, v/v) and analyzed by Hydrophilic Interaction Liquid Chromatography coupled to tandem mass spectrometry (HILIC - MS/MS). Individual metabolites were processed using the Agilent Quantitative analysis software (version B.07.00, MassHunter Agilent Technologies). Relative quantification of metabolites was based on EIC (Extracted Ion Chromatogram) areas for the monitored MRM transitions or accurate masses. The obtained tables (containing peak areas of detected metabolites across all samples) were exported to “R” software http://cran.r-project.org/ and signal intensity drift correction and noise filtering (if necessary, using CV (QC features) > 30%) was done within the MRM PROBS software.

### Image quantification and statistical analysis

Images were analyzed with Fiji software. For manual quantification, the image was duplicated in two windows, one with all fluorescent channels to identify MuSCs and define specific region corresponding to MuSC cytosolic space. These regions were applied to the second window with the brightfield channel for enzymatic activity staining, using the synchronize windows tool. Before quantification, brightfield images were inverted and an unstained region from the same slide was defined as the blank and subtracted to all values from the associated sections. Unless stated otherwise, we used mean values from the quantified region. On dot plots for ISE quantification, each small, light dot represents a single MuSC or fiber, while each large, dark dot of the same color represents the mean for one sample. On correlation each dot represents a single MuSC or fiber, and all dots with the same color are from the same sample. All data are presented as mean of values of independent samples ± s.e.m. Raw data from each independent experiment were analyzed using either an unpaired two-tailed Student’s t-test or a one-way ANOVA. When datasets failed the D’Agostino–Pearson omnibus normality test (α = 0.05), differences were assessed using a two-tailed unpaired nonparametric Mann–Whitney test. Multiple comparisons were corrected by controlling the false discovery rate (FDR < 0.05) using the two-stage step-up method of Benjamini, Krieger, and Yekutieli.

## Author contributions

Conceptualization: AP, PM, CC, MG; Methodology: AP, CC, AB, MQ; Investigation: AP, CC; Supervision: AP, MG; Writing original draft: AP, CC; Writing review & editing: AP, CC, OMD, LAN, MG, PM, FR.

## Data availability

The RNA-seq datasets generated in this study will be deposited in the ArrayExpress database and will be made publicly available upon acceptance of the manuscript (accession number: E-MTAB-17013). All other data supporting the findings of this study are available from the corresponding author upon reasonable request.

## Acknowledgements

The authors thank members of the PFMU, Bioimaging, Histology and ChiRO core facilities (University of Geneva, Switzerland) for experimental support. We thank A. Guguin and A. Henry of the flow cytometry platform of IMRB, Inserm U955, Creteil, France. We thank Prof. M. Rüegg (University of Basel, Switzerland) for the generous gift of TSCmKO mice. We are also grateful to Perrine Castets (University of Geneva, Switzerland) for hosting part of the experimental work and for sharing her expertise on denervation and the TSCmKO model. We acknowledge the LBFA Imaging and cytometry facility (GIS IBiSA, ISdV, LBFA), member of the national infrastructure France-BioImaging supported by the French National Research Agency (ANR-10-INBS-04).

---

**Supplemental Figure 1.**
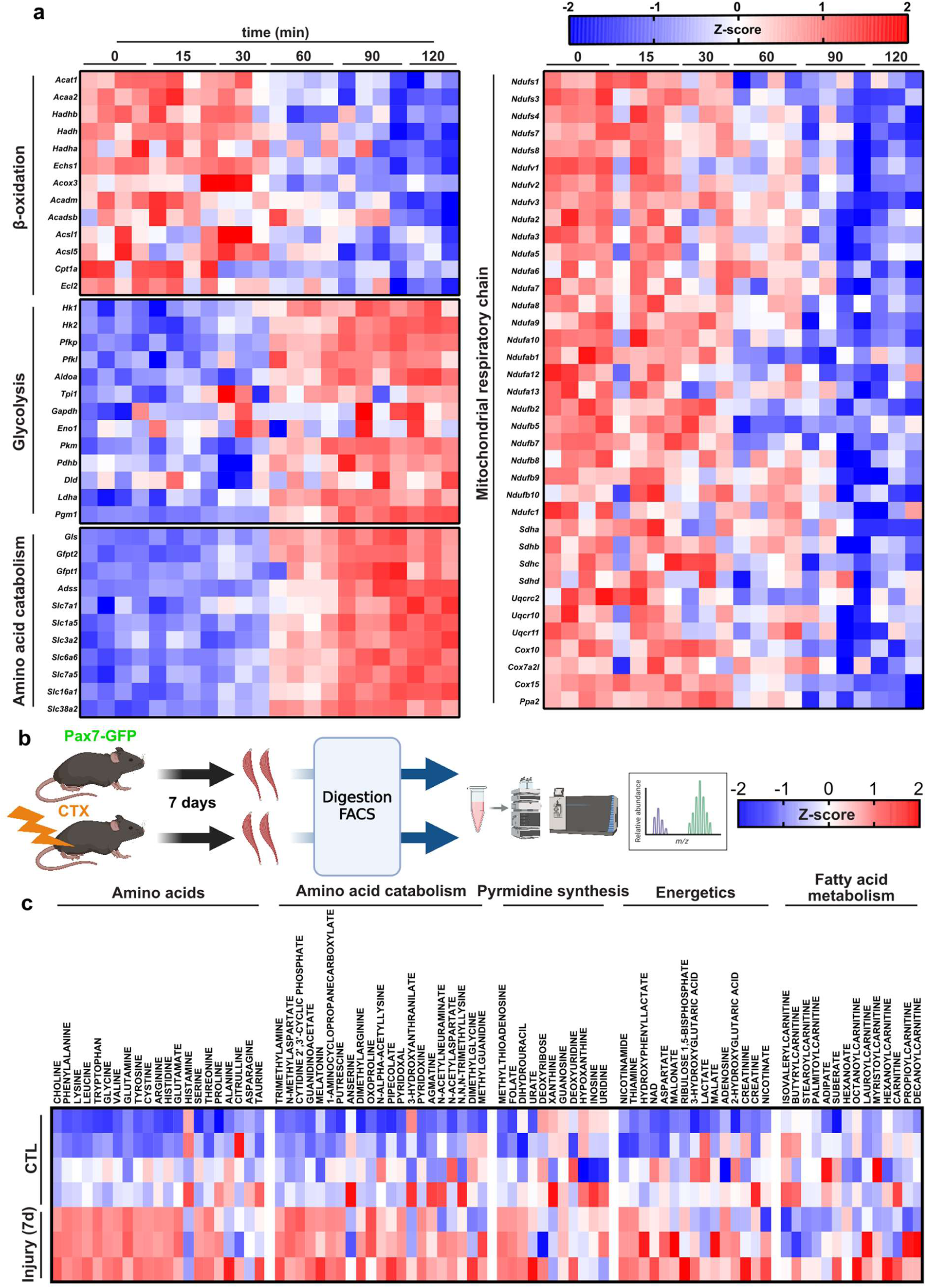
MuSC isolation prevents the characterization of their native metabolism. **a**, Heatmaps showing the expression dynamics of genes involved in fatty acid oxidation, glycolysis, and amino acid catabolism from MuSCs fixed at 0, 15, 30, 60, 90, and 120 minutes after the start of tissue dissociation. **b,** Experimental scheme for isolating unfixed MuSCs (nGFP^+^) by FACS from control or 7 dpi injured muscles for metabolomic analysis. **c,** Heatmap showing the relative abundance of 82 metabolites (columns) in MuSCs isolated from control or injured muscles. Each row represents an individual biological sample.

**Supplemental figure 2.**
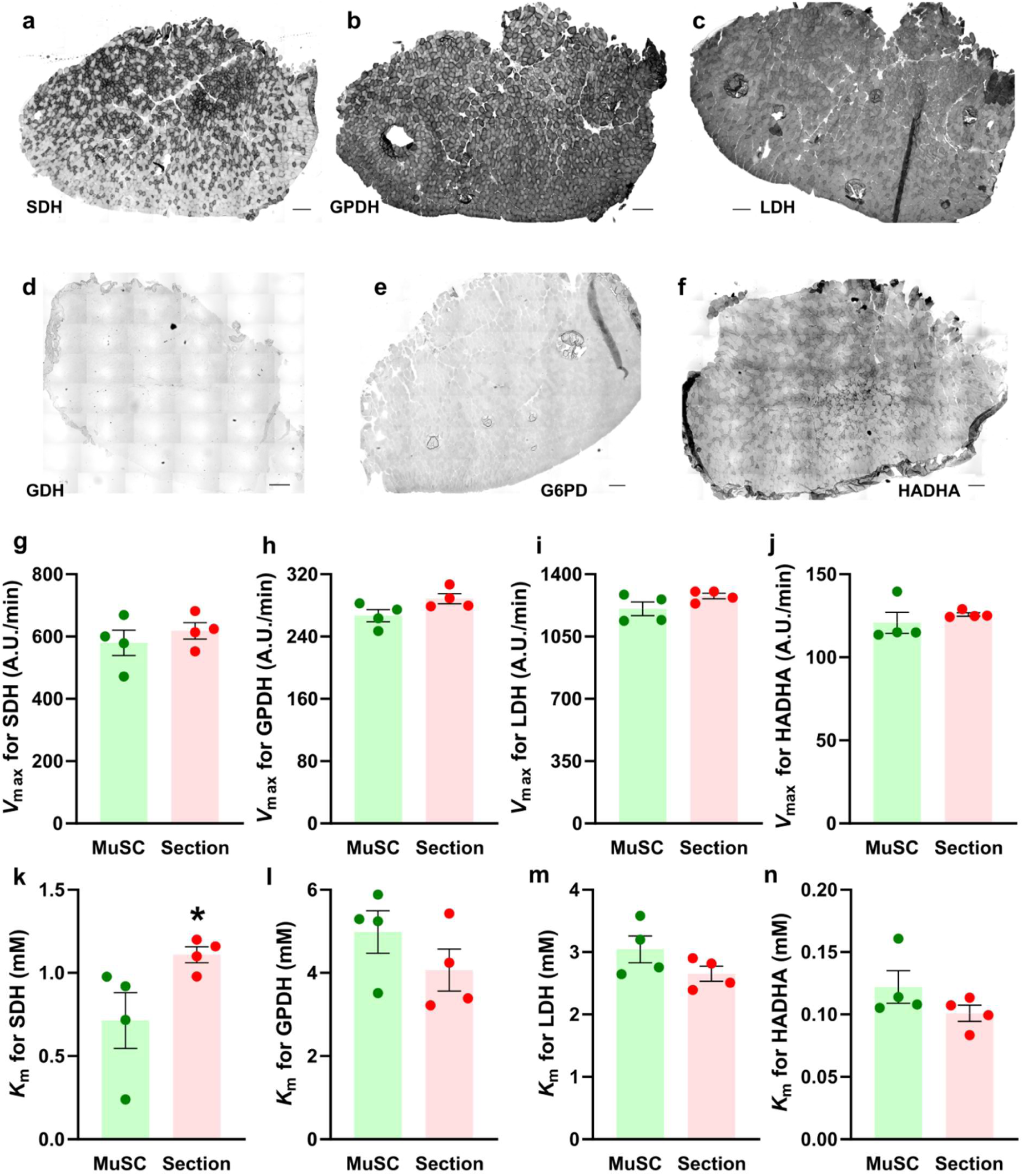
*In situ* enzymatic staining for metabolic profiling of MuSCs., Representative enzymatic staining of the entire *tibialis anterior* sections for SDH **(a)**, GPDH **(b)**, LDH **(c)**, GDH **(d)**, G6PD **(e)** and HADHA **(f)** activities. Scale bars: 200 µm. **g-n,** *V*_max_ (**g-j**) and *K*_m_ (**k-n**) values for SDH (**g, k**), GPDH (**h, l**), LDH (**i, m**) and HADHA (**j, n**) derived from Michaelis-Menten fits of MuSC (green) or whole-section (red) quantifications. *: *p* < 0.05.

**Supplemental figure 3.**
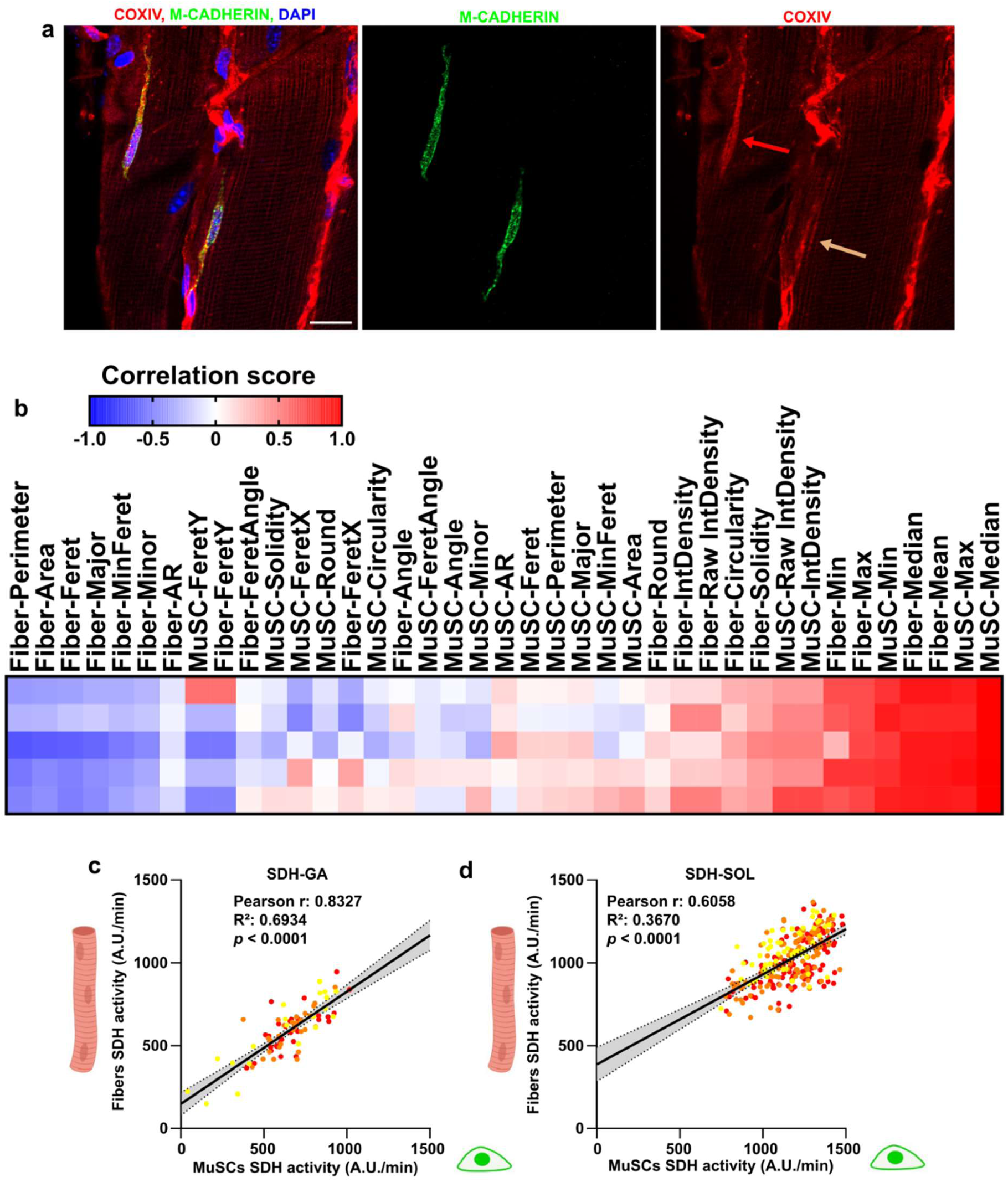
MuSC metabolic heterogeneity and correlation between MuSC and myofiber metabolism. **a-i**, Immunostaining of COXIV (red, mitochondria) and M-Cadherin (green, MuSCs) showing in right panel an oxidative (red arrow) and a glycolytic (beige arrow) MuSC. Scale bar: 20 µm. **b,** Heatmap showing the correlation score between MuSC SDH mean staining and other parameters in resting *tibialis anterior*, each row represents a different sample. **c-d,** Linear regression analysis of SDH activity in superficial *gastrocnemius* (**c**) or *soleus* (**d**) in fibers (y-axis) and associated MuSCs (x-axis) with Pearson correlation scores, revealing a correlation between the metabolic profiles of fibers and associated MuSCs.

**Supplemental Figure 5.**
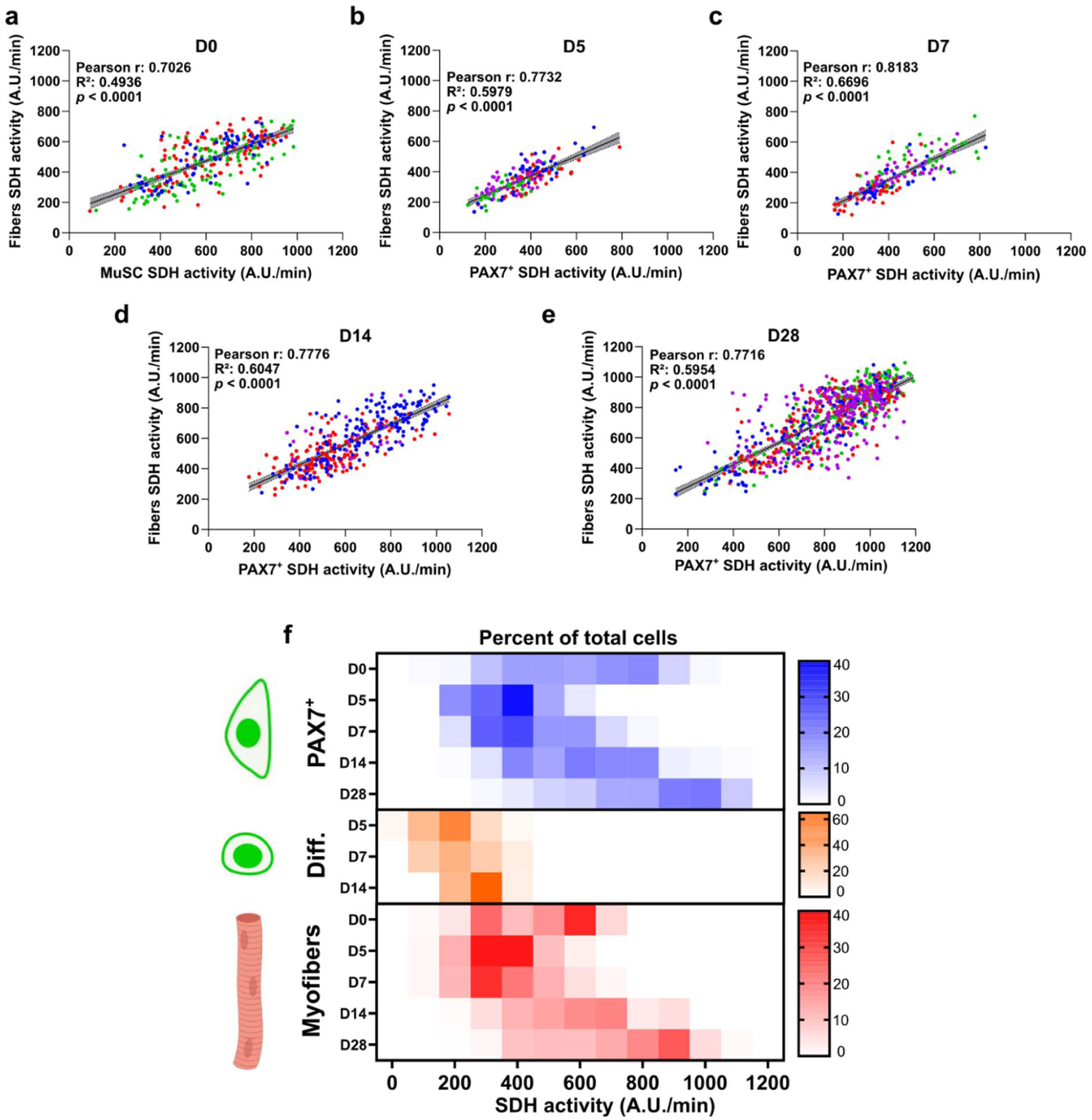
Mapping of oxidative metabolism during muscle regeneration. **a–e**, Linear regression analyses of SDH activity in *tibialis anterior* Xfibers (y-axis) and associated PAX7^+^ cells (x-axis), with Pearson correlation scores shown for D0 (**a**), D5 (**b**), D7 (**c**), D14 (**d**), and D28 (**e**) after injury. **f**, Heatmap showing the distribution of cells (as % of the total population) according to their SDH activity clustering in PAX7^+^ (blue), Differentiated cells (orange), and myofibers (red).

